# Functional gradients reveal altered functional segregation in patients with amnestic mild cognitive impairment and Alzheimer’s disease

**DOI:** 10.1101/2022.11.16.516698

**Authors:** Yirong He, Qiongling Li, Zhenrong Fu, Debin Zeng, Ying Han, Shuyu Li

## Abstract

**Background:** Alzheimer’s disease (AD) and amnestic mild cognitive impairment (aMCI) are associated with disrupted functional organization in brain networks, yet the finer changes in the topological organization in aMCI and AD remain to be investigated. Connectome gradients are a new tool representing brain functional topological organization in a low-dimensional space to smoothly capture the human macroscale hierarchy.

**Methods:** Here, we examined altered topological organization in aMCI and AD by connectome gradient mapping. We further quantified functional segregation by gradient dispersion. Then, we systematically compared the alterations observed in aMCI and AD patients with those in normal controls (NCs) in a two-dimensional functional gradient space from both the whole-brain level and module level.

**Results:** Compared with NCs, the first gradient, which described the neocortical hierarchy from unimodal to transmodal regions, showed a distributed and significant suppression in AD patients, while abnormalities were only limited to local regions in aMCI patients. The second gradient showed a decreased pattern in the somatomotor module in both aMCI and AD patients. Furthermore, gradient dispersion showed significant decreases in AD patients at both the global level and module level, whereas this alteration was limited only to limbic areas in aMCI. Notably, we demonstrated that suppressed gradient dispersion in aMCI and AD patients was associated with cognitive scores.

**Conclusions:** Changes in functional gradients could reflect different degrees of altered brain network segregation in aMCI and AD. These findings provide new evidence for altered brain hierarchy in aMCI and AD, which strengthens our understanding of the progressive mechanism of cognitive decline.

## 1 Background

Alzheimer’s disease (AD) is the most common type of dementia that often occurs in middle-aged and elderly people, is characterized by devastating neurodegeneration, and is accompanied by memory loss and other declines in cognitive function (1, 2). Patients with amnestic mild cognitive impairment (aMCI) are at high risk for developing AD and present with memory impairment, but aMCI does not significantly interfere with daily functioning (3). Neuroimaging-based methods have revealed abnormities in patients with aMCI and AD, such as decreased gray matter volume, atrophic hippocampus, and damaged white matter fiber tracts (4-6). Connectome-based approaches have revealed disrupted topological organization in large-scale brain networks, such as disrupted small-world architecture, and disconnections between brain regions (7, 8).

Brain networks are organized in a hierarchical modular architecture (9). Modular brain networks are patterns of functional segregation characterized by a balance of dense connections within modules and sparse connections between modules (10, 11). The position of each brain region in the hierarchical organization corresponds to its specific functional role, and it allows integration among networks and the segregation of specialized functional domains (12-14). The altered functional roles in the hierarchical organization may attribute to abnormal functional segregation. Although numerous graph theory-based studies have reported abnormal functional network changes in aMCI and AD, the inconsistent results regarding functional segregation suggest that this issue still needs to be further investigated (15). More importantly, most previous studies examined functional segregation by computing local functional characteristics based on functional connection strength and captured coarse-grained representation (15). While abnormalities in diseased brain networks are not confined to the local functional system, the damage can propagate throughout the brain network, especially in critical regions, which may result in more serious disturbances such as global alterations and functional brain network reconfiguration (16, 17). From the perspective of the hierarchical organization of functional brain network reconfiguration, it is still unclear what global functional segregation patterns that aMCI and AD follow. Corresponding to hierarchy, a large-scale organization axis named the “sensory-fugal axis” provides a framework for describing and ordering the cortical regions globally and can be used to investigate changes in aMCI and AD in a more detailed manner (12).

Recently, nonlinear dimensionality reduction approaches have offered the chance to map high-dimensional functional connectivity data into a low-dimensional space, such as diffusion map embedding (13, 18). This approach revealed multidimensional functional organization axes along which the functional connection profiles are smoothly variable in the neocortex. The components of functional connectivity in the low-dimensional space were identified as functional connectome gradients and explain connectome variances in descending order. The first gradient, corresponding to the “sensory-fugal” axis, reveals the macroscale functional hierarchy of functional connectivity profiles from primary to transmodal regions (12, 13). The second gradient dissociates the visual and somatomotor regions (13). By continuously ordering the regions according to their functional connectivity profiles on the gradient axes, the position of each region in the gradient space can be regarded as having a unique functional role in neocortical organization (13, 19-21). Connectome gradients have been applied for the characterization of brain organization features to investigate multiple critical topics, such as cognition-related terms (22), the relationship between structure and function (23, 24), brain development (25), brain aging (26), and neurological and psychological disorders (23, 27-30). This approach can be used to study the reconfiguration of the functional network related to aMCI and AD by carefully arranging brain regions in a continuous gradient space according to the functional connectivity profiles. This approach may offer a more accurate description of cortical regions to reveal the abnormalities of functional network segregation in aMCI and AD.

To investigate alterations in functional segregation in aMCI and AD, the present study applied connectome gradient analysis based on resting-state functional magnetic resonance imaging (rs-fMRI) datasets collected from three groups: normal controls (NCs) (n = 42), aMCI patients (n = 43) and AD patients (n = 41). By mapping functional connectivity into a low-dimensional continuous gradient space, we investigated altered segregation in patients with aMCI and AD. We hypothesized that the disease-related changes in the neocortex would cause reconfiguration of the functional network and further influence functional segregation.

## 2 Methods

### 2.1 Participants

In this study, we recruited 43 aMCI patients and 41 AD patients from the memory clinic of the Neurology Department of Xuanwu Hospital, Capital Medical University, China, and 42 normal controls were enrolled from local communities in Beijing, China. This study was approved by the Ethics of the Medical Research Ethics Committee in Xuanwu Hospital, and written informed consent was obtained from all participants. All participants underwent standard clinical assessments, including a medical history investigation, neurological examination, and neuropsychological tests. The cognitive tests included the Montreal Cognitive Assessment (MoCA, Beijing version) (31), Auditory Verbal Learning Test (AVLT), Clinical Dementia Rating (CDR) (32), Hamilton Depression Rating Scale (HAMD), Activities of Daily Living (ADL) scale, Hachinski Ischemic Scale and the Center for Epidemiologic Studies Depression scale (33).

As described in our previous study (34), the diagnostic criteria for aMCI patients followed the Petersen criteria (3), which were defined as follows: (1) memory complaints, confirmed by an informant; (2) significant decline in objective memory measured by the MoCA adjusted for years of education and AVLT; (3) CDR score = 0.5; and (4) the exclusion of subjects with other subtypes of MCI. AD patients were diagnosed based on the criteria of the National Institute of Aging-Alzheimer’s Association (NIA-AA) for dementia due to AD (35), including: (1) symptoms that conformed with the diagnostic criteria for dementia; (2) brain atrophy in the hippocampus based on structural MRI; (3) symptoms that lasted for more than 6 months; and (4) CDR score≥ 1.

The exclusion criteria for this study were as follows: (1) major psychiatric diagnosis; (2) an abnormal Hachinski Ischemic Scale; (3) left-handedness; (4) impaired executional, visual, or auditory functions; (5) other types of neurological disease that may hinder the project; (6) history of serious alcohol or drug abuse; and (7) images with large head motion.

### 2.2 MRI acquisition and processing

All images were acquired on 3.0 T Siemens system (Magnetom Trio Tim; Erlangen, Germany) at the Department of Radiology, Xuanwu Hospital, Capital Medical University, Beijing, China. T1-weighted images were acquired using a magnetization-prepared rapid gradient echo (MPRAGE) sequence (TR = 1900 ms; TE = 2.2 ms;TI = 900 ms; flip angle = 9^°^; FOV = 224 mm × 256 mm ; matrix size = 448 × 512 ; number of slices = 176 ; slice thickness = 1 mm). Functional images were acquired axially using a gradient-echo EPI sequence (TR = 2000 ms; TE = 40 ms ; flip angle = 90^°^ ; FOV = 240 mm × 240mm ; matrix size = 64 × 64 ; number of sections = 28 ; section thickness = 4 mm ; voxel size = 3.75 × 3.75 × 4mm^3^ ; gap = 1 mm; volume number = 239).

For fMRI preprocessing, we used the FMRIPREP 20.0.7 pipeline (36). T1w images were preprocessed as follows: intensity nonuniformity correction, skull-stripping, spatial normalization to standard space (MNI152NLin6Asym), brain tissue segmentation, and reconstruction of cortical surfaces. The steps of rs-fMRI data preprocessing were as follows: removing the first five timepoints, motion correction and slice-timing correction, and coregistration to the corresponding T1w image using boundary-based registration. Then, we performed spatial smoothing using an isotropic Gaussian kernel of 6 mm full-width at half-maximum. Following a denoising step, we performed ICA-based Automatic Removal Of Motion Artifacts (AROMA) and linear regression, including the averaged white matter and cerebrospinal fluid signals as nuisances (37, 38). Then, we applied high-pass filtering using a discrete cosine filter with 128s cut-off. The preprocessed time series in volume space were then sampled at each vertex on cortical surfaces and registered to the fsaverage5 surface template (10k vertices per hemisphere). Many internal operations of FMRIPREP use Nilearn, principally within the BOLD-processing workflow. For more details of the pipeline see https://fmriprep.readthedocs.io/en/stable/workflows.html.

### 2.3 Functional connectome and gradient analysis

In this study, we generated a functional connectivity network for each subject by computing the pairwise Pearson’s correlation (with 20484 × 20484 entries) between the time series of all vertices. According to previous studies, we thresholded this connectivity matrix to reserve the top 10% of edges per row (13, 27). Then, a normalized angle affinity matrix was obtained by computing the row-wise similarity of functional connectivity profiles (26, 39). The following steps in the BrainSpace Toolbox were used to compute the functional gradients (40). The normalized angle matrix was then fed into the diffusion map embedding algorithm, which was a nonlinear dimensionality reduction approach (18). In line with previous studies, we set parameter *α* = 0.5 to retain the global relations between data points in the embedding space (13, 27, 41). To make the gradients comparable across individuals and eliminate the randomness of the direction of the gradients, we performed Procrustes rotations to align the individual gradients to a well-established group-level cortical gradient map derived from the Human Connectome Project (HCP) dataset (13, 42). For each group, we generated a group-level functional connectivity matrix by averaging the individual functional connectivity matrices and applied the same procedures as above, thus obtaining group-level gradients.

### 2.4 Derivation of the group-level network modular structure

According to previous studies, the modular structure in healthy elderly individuals was different from those in young adults, as evidenced by a less distinct modular structure, decreased segregation, and functional specialization (43, 44). In this study, instead of using a prior network partition based on young healthy adults, we generated a group-level network modular structure based on NCs to describe a more accurate network partition for elderly individuals. Given the different numbers of modules among individuals, this procedure was performed at the group-level functional network. Based on the mean functional connectivity network of the NC group, we performed community detection using Louvain methods to generate a group-level modular structure (45). In this procedure, the spatial resolution parameter γ determines the number and size of the modules. We set *γ* = 1.12 to produce a similar number of modules to those previously reported (46, 47). Considering the randomness of Louvain community detection, we repeated this algorithm 1000 times. Then, we averaged the 1000 modular structure results and obtained a stable modular structure at the group-level. We computed the mean normalized mutual information (NMI) to measure the similarities between 1000 partitions (48).

### 2.5 Gradient dispersion analysis

To depict the multidimensional features of cortical functional organization, we used the concept of “gradient dispersion” from previous research and modified it (26). This modified concept was used to quantify the dissimilarity of functional connectivity profiles between brain regions. In this study, the first two gradients were used to construct a 2D gradient space. For each subject, the global dispersion was calculated as the sum of Euclidean distances of each node to the centroid in the 2D gradient space. To describe the dissimilarity of functional connectivity features between one module and other modules, between-module dispersion was quantified as the mean Euclidean distance from the centroid of a module to the centroids of all the other modules.

### 2.6 Graph-theoretic analysis

To explore the relationship between gradient dispersion and functional specialization, we assessed how gradient dispersion was related to connectome topology. We calculated the graph-theoretic system segregation (49) and used the Brain Connectivity Toolbox (https://sites.google.com/site/bctnet/) to compute the participation coefficient (50, 51). For each subject, consistent with the gradient analysis, we used the same threshold matrix as the input. System segregation was used to characterize functional specialization, containing within- and between-modular functional connectivity (49). It was defined as follows:

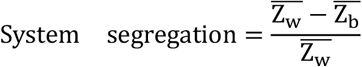

We computed the system segregation for each functional module and the whole brain. At the module level, 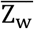 is the mean connectivity between all paired nodes within the given module, whereas 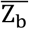 is the mean connectivity between all nodes within the given module and nodes outside of that module. At the whole-brain level, 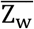 is the mean connectivity between all paired nodes within the same modules, and 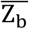 is the mean connectivity between all paired nodes in different modules (49).

In addition, the participation coefficient was used to measure inter-module connectivity and quantify the extent of a node participating in other modules. We computed the mean participation coefficient across all nodes from a given functional module at the module level and averaged it across the whole brain (50, 51).

### 2.7 Statistical analysis

We performed analysis of variance (ANOVA) to compare the group differences in age, years of education, and neuropsychological test scores, including MoCA and AVLT scores. The chi-square test was performed to compare sex differences between groups.

In this study, we focused on the first two gradients, which explain the largest two variances. We compared the gradient values of the aMCI group and AD group with those of the NC group at both the macroscale functional module level and vertex level. For the comparisons at the module level, we computed the mean values of the gradients for each module. Furthermore, we statistically compared the global dispersion and between-module dispersion.

The statistical analysis for the above measures between the three groups was conducted using general linear models in SurfStat (http://www.math.mcgill.ca/keith/surfstat/), controlling for age, sex, years of education and head motion (mean frame-wise displacement, mFD). Surface-based results were multiple comparison corrected for family-wise errors (FWE) using a random field theory (*p*_*FWE*_ < 0.05). Module-based results were corrected with the false discovery rate (q_FDR_ < 0.05).

### 2.8 Relationships between gradient dispersion and cognitive scores

To test the hypothesis that decreased brain functional segregation is related to decreased cognitive performance in patients with cognitive disease, we used gradient dispersion to represent functional segregation to explore this relationship in aMCI and AD groups. We chose MoCA, AVLT immediate recall, AVLT delayed recall, and AVLT recognition scores to quantify the cognitive performance of patients. Then, we performed linear regressions to explore the relationship between individual cognitive performance and global dispersion/between-module dispersion of each module.

## 3 Results

### 3.1 Cognitive performance and demographic data comparisons

Compared with the NC group, there were no significant differences in age, sex, or years of education in the aMCI group or the AD group (*p* > 0.05). Moreover, the neuropsychological test scores (MoCA, AVLT immediate recall, AVLT delayed recall, and AVLT recognition) were significantly lower in the aMCI group and the AD group than in the NC group (*p* < 0.001). The results are shown in Table 1.

**Table 1.** Cognitive performance and demographic data.

|  | NC | aMCI | AD |
| --- | --- | --- | --- |
| **Number** | 42 | 43 | 41 |
| **Age (y)** | 64.24±6.16 | 67.47±10.03 | 68.88±7.86 |
| **Sex (Male:Female)** | 15:27 | 21:22 | 17:24 |
| **Years of education** | 11.17±5.61 | 10.44±4.96 | 9.68±4.71 |
| **MoCA** | 26.02±2.95 | 19.67±4.28* | 13.10±5.46* |
| **AVLT immediate recall** | 9.32±1.94 | 5.84±1.34* | 3.53±1.58* |
| **AVLT delayed recall** | 10.43±2.31 | 3.19±2.81* | 0.98±1.60* |
| **AVLT recognition** | 12.07±2.13 | 6.58±4.28* | 3.51±3.08* |

NC, normal control; aMCI, amnesic mild cognitive impairment; AD, Alzheimer's disease. MoCA, Montreal Cognitive Assessment (Beijing version). AVLT, auditory verbal learning test. \*Represents a significant difference between the NC group and other groups ( $p < 0.001$ ).

### 3.2 Group-level modular structure of the NC group

Based on the group-level functional connectivity matrix of the NC group, we identified 6 modules and then labeled them according to the network definition of previous studies (46, 47): somatomotor network (SN), visual network (VN), ventral attention network (VAN), dorsal attention network (DAN), limbic network (LN), and default mode network (DMN) (Fig. 1b). The mean NMI was 0.9791 ± 0.0465 between 1000 partitions, which indicated that our partition was highly reliable.

**Fig. 1.**
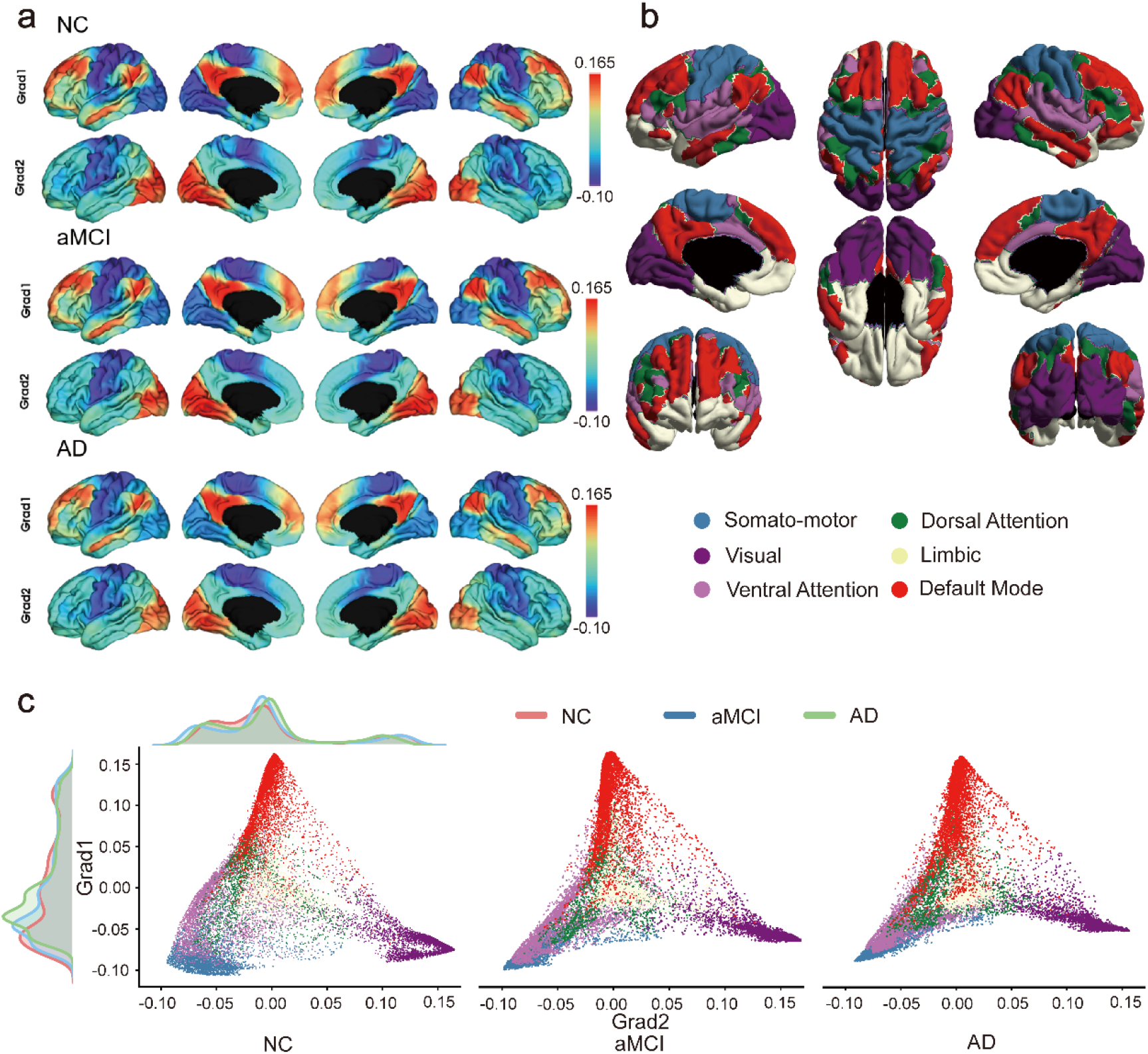
Functional gradient mapping in the NC, aMCI, and AD groups. **a** The first two gradients in each group. From top to bottom are the NC, aMCI and AD groups. **b** Modular structure of elderly participants derived from the NC group-level connectome. Labels were assigned based on networks defined in (46, 47). **c** The first two functional gradients mapped into a 2D gradient space and colored by modules. From left to right are the NC, aMCI, and AD groups. The density map shows the gradient distributions of the NC (red), aMCI (blue) and AD (green) groups. The y-axis denotes the first gradient, which represents the transition from unimodal regions to transmodal regions. The x-axis denotes the second gradient, which separates the visual network and somatomotor network. Normal controls, NCs; amnesic mild cognitive impairment, aMCI; Alzheimer’s disease, AD; grad, gradient.

### 3.3 Macroscale gradients in the NC, aMCI, and AD groups

By applying the diffusion map embedding method, we obtained a set of components (13). We included the first two gradients in this study, which accounted for the largest variance. The first gradient explained 23 ± 4% of variance in the NC group, 23 ± 5% in the aMCI group, and 20 ± 5% in the AD group, which showed a significant decrease compared with the NC group (*p* < 0.05), indicating that the dominant role of the first gradient in the AD group was weakened. The second gradient explained 14 ± 1% of variance in the NC group, 15 ± 2% in the aMCI group and 15 ± 2% in the AD group (Supplementary Materials Fig. 1[see Additional file 1]). Consistent with previous studies (13) (26), the first gradient reflected the transition from unimodal to transmodal areas. The second gradient separated the VN and SN. The overall gradient patterns were similar in the NC, aMCI, and AD groups but with slight changes (Fig. 1a). We applied our modular structure to label vertices in the neocortex and were represented in the 2D gradient space (Fig. 1c). Globally, the aMCI group and AD group, especially the AD group, which had a more abnormal distribution, showed altered gradient distributions compared with the NC group. Specifically, the first two gradients of the AD group were contracted compared with those of the NC group and aMCI group, whereas there was a slight change in the aMCI group compared with the NC group.

### 3.4 Gradient alterations at the vertex and module levels

We compared the gradient alterations of the aMCI and AD groups with those of the NC group at the vertex level and module level. Vertex level results showed that the aMCI group and AD group had similar alteration patterns in the two gradients, while the AD group had a more serious degree of alteration. After multiple comparison correction for family-wise errors (*p*_*FWE*_ < 0.05), we found that transmodal areas, including the left angular gyrus, decreased, whereas unimodal areas, such as the visual area and somatomotor area, increased in the first gradient of the AD group. For the second gradient, there was only a slight decrease in the somatomotor area in the AD group (Fig. 2a). We applied our modular structure to label the vertices on the neocortex to summarize our findings of gradient alterations at the module level. Similar to the findings at the vertex level, the results showed that the VAN and DMN significantly decreased, whereas the SN and VN significantly increased for the first gradient in the AD group (false discovery rate q_FDR_ < 0.05). There was a marginal decrease in VAN for the first gradient in the aMCI group (*p*_*uncorrected*_ < 0.05). For the second gradient, SN showed a slight decrease in both the aMCI and AD groups (*p*_*uncorrected*_ < 0.05) (Fig. 2b).

**Fig. 2.**
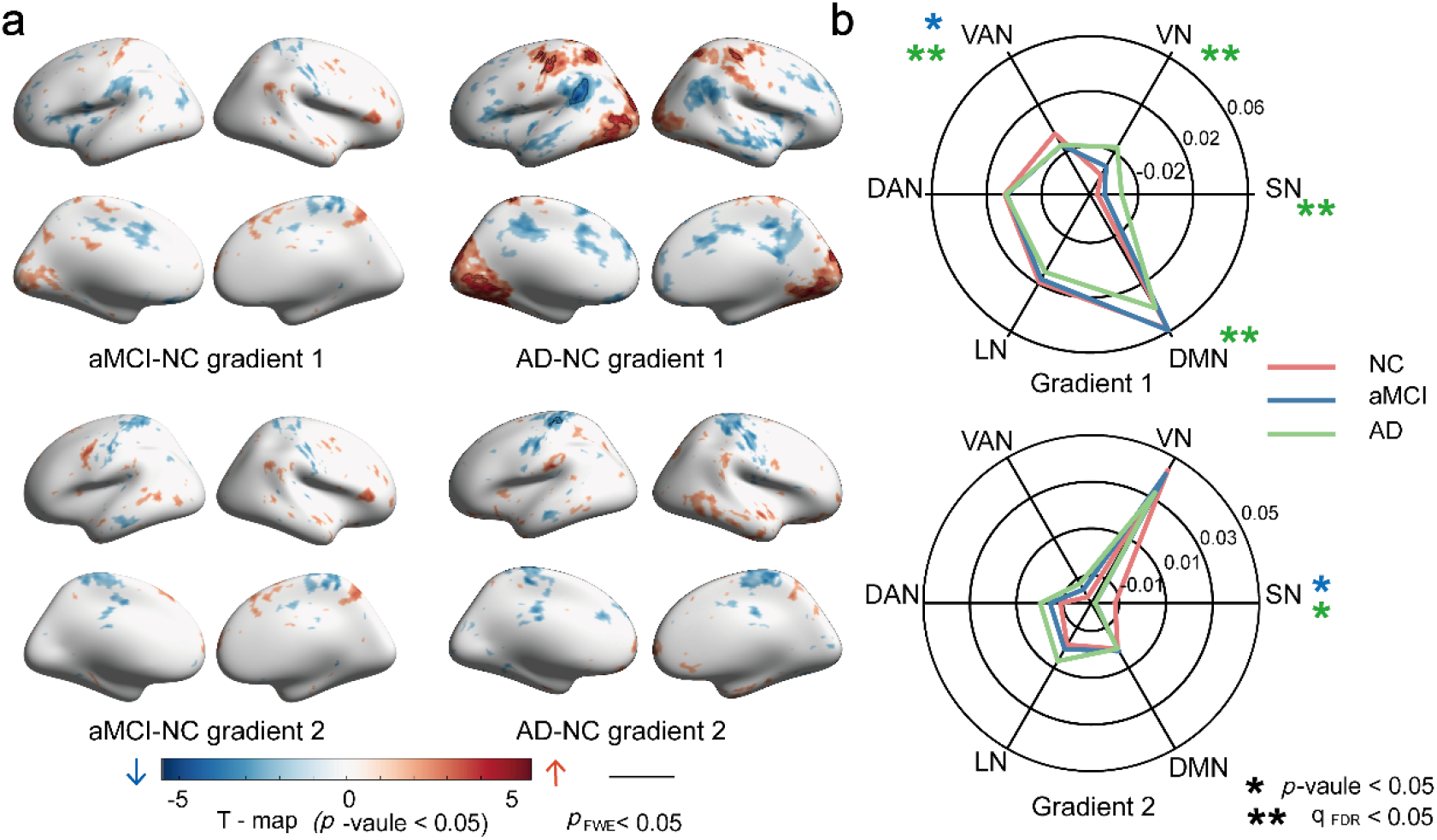
Functional gradient comparisons at both the vertex level and the module level. **a** Vertex-based statistical comparisons between groups. Increases/decreases in aMCI/AD are shown in red/blue, and results surviving family-wise error (FWE) correction are plotted as black lines. **b** Module-based statistical comparisons between NC (red) and aMCI (blue) and AD (green). The radar plot shows the mean within-module gradients of the three groups. *Represents a significant difference between the NC group and other groups (*p* < 0.05). **Represents the results corrected for false discovery rate (q_FDR_ < 0.05).

### 3.5 Differences in gradient dispersion in the NC, aMCI, and AD groups

As shown in Fig. 1c, each module occupied a position in the 2D gradient space. Consistent with previous studies (13, 26, 27), the SN, VN, and DMN occupied the extreme positions of the triangle, and other networks were distributed in the middle of the space. The nodes from the same module were close to each other, whereas nodes from different modules were far apart. Global gradient dispersion was used to measure the segregation degree of the whole brain. Comparisons of global dispersion between diagnostic groups revealed a significant decrease in the AD group versus the NC group (*p* < 0.01). There was no significant difference between the aMCI group and NC group (Fig. 3a). We adopted two previously established functional topological measures related to brain functional specialization, system segregation and the participation coefficient, to investigate the correlation with global dispersion (49-51). System segregation showed a significant positive correlation with global dispersion (r = 0.3734, *p* < 0.001), whereas the participation coefficient was significantly negatively correlated with global dispersion (r = −0.4620, *p* < 0.001) (Fig. 3b). This result suggested that the decreased global gradient dispersion in the AD group reflected decreased brain network segregation.

**Fig. 3.**
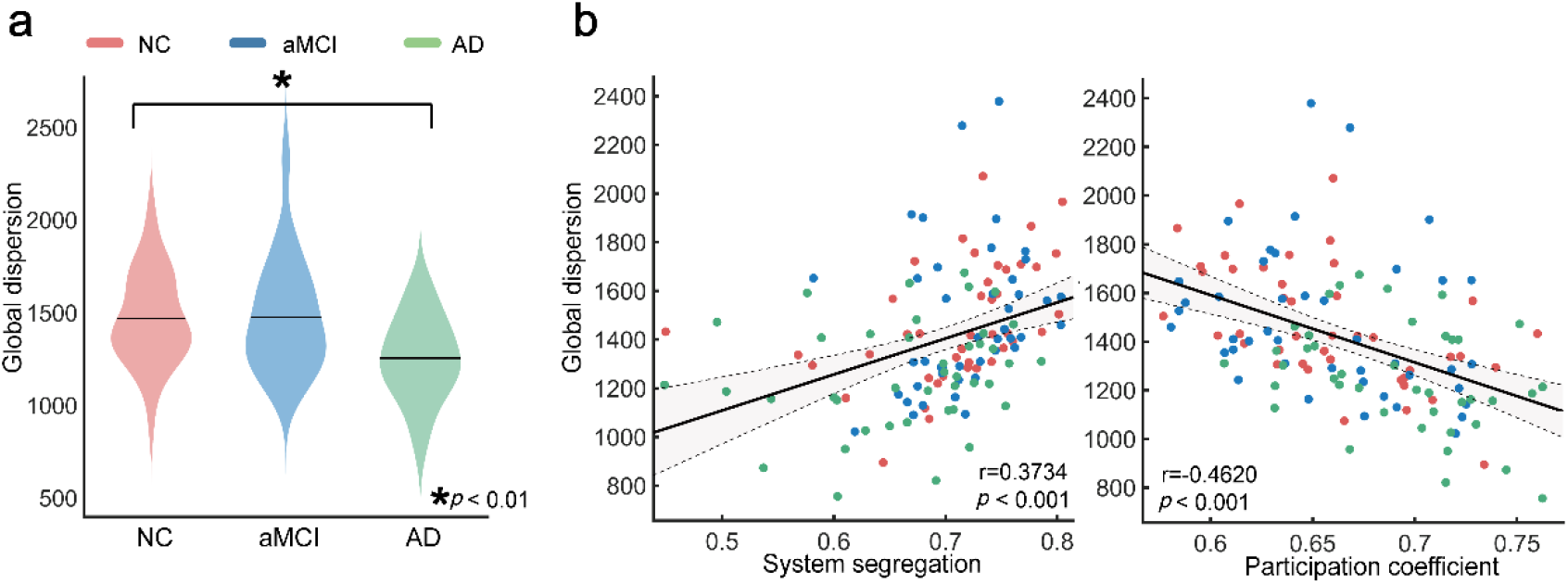
Global dispersion analysis. **a** Global dispersion comparisons between the NC (red) and aMCI (blue)/AD (green) groups. There was a significant reduction in the AD group compared with the NC group. **b** Relationships between global dispersion and system segregation or the participation coefficient. Global dispersion is positively correlated with system segregation (r = 0.3734, *p* < 0.001) and negatively correlated with the participation coefficient (r = −0.4620, *p* < 0.001).

We further compared the between-module dispersion for each module. This measure estimates the dissimilarity between the given module functional role and other modules. According to the results of the between-module dispersion shown in Fig. 4a, most of the modules except for LN in the AD group showed significantly decreased between-module dispersion compared to the NC group (q_FDR_ < 0.05), whereas the aMCI group showed an increased pattern limited only in LN (*p*_*uncorrected*_ < 0.05). Similar to the global dispersion analysis, we used the t-statistic to measure differences between the NC group and aMCI/AD groups in between-module dispersion, system segregation, and the participation coefficient. As shown in Fig. 4b, in the AD group, the changes in between-module dispersion and system segregation changes were in the same direction and opposite to the participation coefficient. In the aMCI group, the absolute value of the t-statistic of between-module dispersion was larger than the other two measures in most of the modules, indicating that gradient dispersion was more sensitive in a diseased state.

**Fig. 4.**
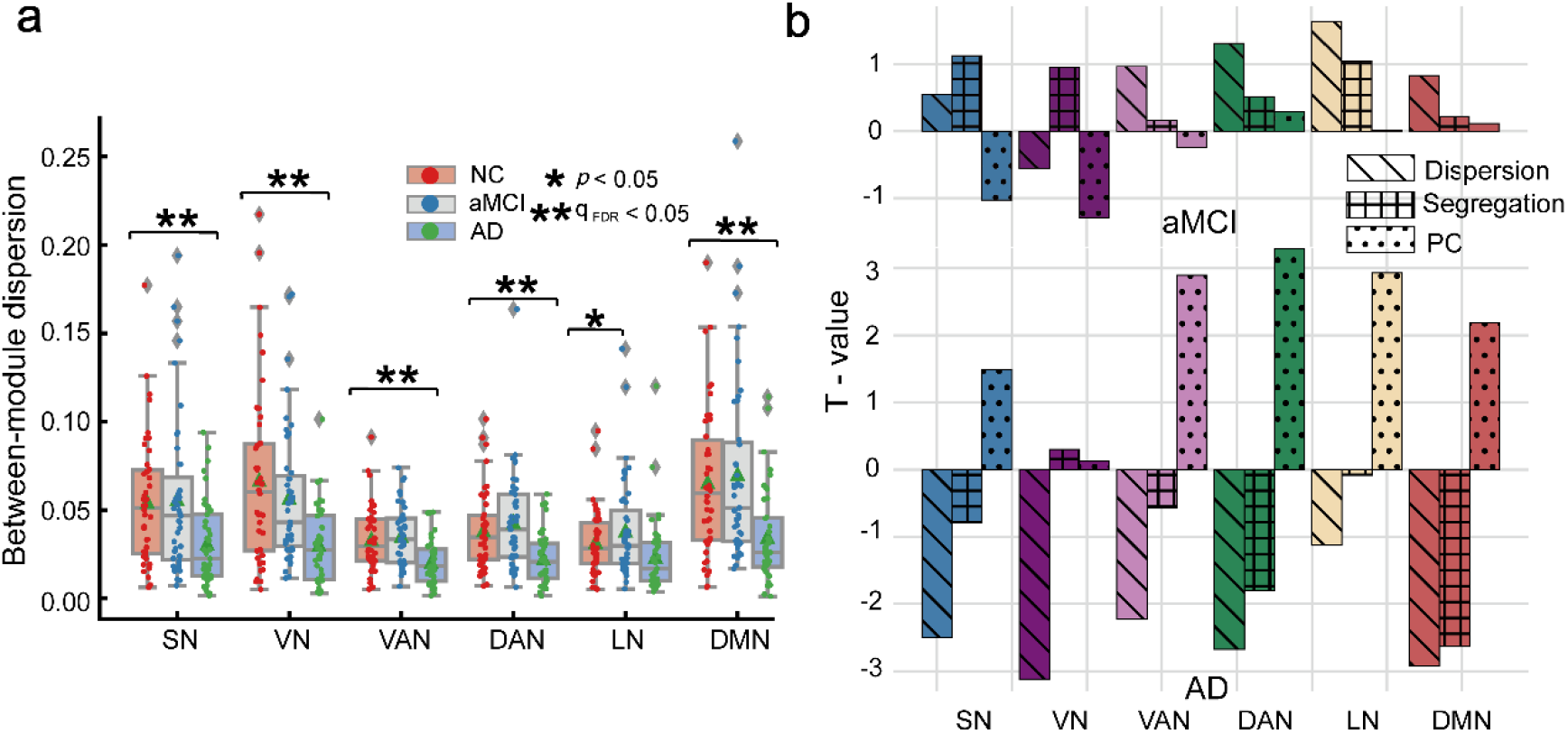
Between-module dispersion analysis. **a** Between-module dispersion comparison between the NC and aMCI/AD groups. The results showed a significant reduction in the somatomotor network (SN), visual network (VN), ventral attention network (VAN), dorsal attention network (DAN), and default mode network (DMN) in AD patients compared with NCs (q_FDR_ < 0.05) and an increase in the limbic network (LN) in aMCI patients compared with NCs (*p*_*uncorrected*_ < 0.05). **b** T-statistic differences in three groups for between-module dispersion, system segregation, and the participation coefficient (PC). The top Figure represents aMCI-NC, and the bottom Figure represents AD-NC. In general, the absolute value of the t-statistic of between-module dispersion was larger than the other two measures.

### 3.6 Relationships between gradient dispersion and cognitive scores

We performed linear regressions between individual cognitive scores and global dispersion/between-module dispersion. In this study, the MoCA and AVLT were adopted to measure cognitive performance. The results showed that dispersion was positively correlated with the cognitive scores at both the global and module levels. Except for LN, all the other modules showed a significant correlation with the cognitive scores (Fig. 5). Our findings suggest that gradient dispersion has the potential to be a predictor of cognitive performance in patients with aMCI and AD.

**Fig. 5.**
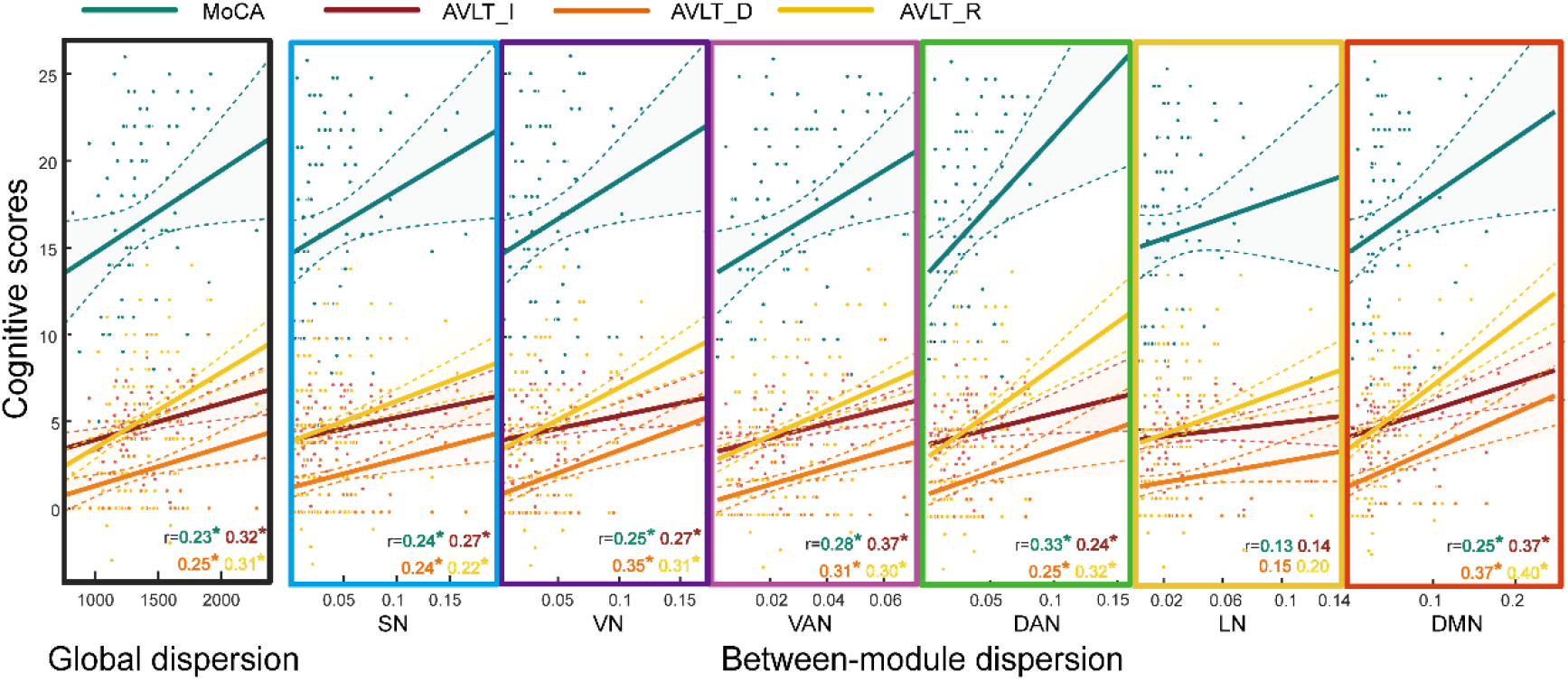
Relationships between dispersion and cognitive scores in aMCI and AD patients. The MoCA (green), AVLT immediate recall (AVLT_I, dark red), AVLT delayed recall (AVLT_D, orange), and AVLT recognition (AVLT_R, yellow) scores are positively correlated to global dispersion and between-module dispersion in 6 modules. Lines representing linear regression between dispersion and cognitive scores are depicted for each scatterplot. *Represents a significant correlation between gradient dispersion and the cognitive score (*p* < 0.05).

## Discussion

In this study, we combined the concepts of hierarchy and modular structure by applying connectome gradient analysis to study abnormal functional segregation in patients with aMCI and AD. We demonstrated that in patients with aMCI and AD, the hierarchical organization remained, which spanned from unimodal to transmodal regions. However, the abnormality degree of the gradients was related to the severity of the disease. By measuring the alterations in functional gradient dispersion, our results revealed abnormal functional segregation in patients with aMCI and AD. There were global changes in patients with AD, whereas altered gradients were limited to local regions in patients with aMCI. Cognitive correlation analysis demonstrated that gradient dispersion can potentially predict cognitive performance. Our findings provide evidence for abnormal functional segregation revealed by functional gradients in patients with aMCI and AD and strengthen the understanding of the functional gradient and progressive mechanism of cognitive decline.

### Functional gradient space

By continuously ordering the cortical regions in a low-dimensional space, gradients have been applied to investigate the macroscale organization of the human brain in a more detailed way (13) (14). The present study constructed a 2D functional gradient space. Based on the concept of “connectivity fingerprints”, we classified the position of each region in the gradient space as its unique role in the functional organization (20, 52). This concept claimed that the functional role of a region is determined by its pattern of connection with the rest regions of the cerebral cortex (20, 52). We described each region using connection patterns rather than the physical location so that we could more accurately characterize the function of each region (14, 42, 53). The results of Fig. 1c show that regions from the same module or modules with similar functions were close to each other in the gradient space. For example, the DAN and VAN were next to each other, while the DMN and sensory networks were far apart because of their very different functions. Previous studies suggested that variations in the range of gradient values represent changes in functional organization (27, 29). A compression in the range of gradient values suggests similarity in the functional roles of modules and a decrease in the functional segregation.

### Altered neocortical hierarchy in patients

Our results showed altered functional gradients in patients with aMCI and AD. The gradients in patients with AD were suppressed and had a more distinguished pattern than those with aMCI, indicating altered functional segregation in patients. Most modules in patients with AD showed abnormal gradient values, which indicated global changes in functional networks in AD. Although the abnormalities of aMCI were slight, the results in a few regions overlapped with those of AD, which suggests a continuous change in the progression of cognitive disease, and abnormalities can spread from local regions to the global brain network. Note that most of the changes were concentrated on the principal gradient, which corresponded to the “sensory to fugal” axis spanning from unimodal to transmodal regions in the cortex (Fig. 2) (12). This axis is regarded as the core axis of the human brain cerebral cortex reflecting a variety of diverse neurobiological features across the cortex, supporting the functional processing hierarchy from sensation and action to more abstract cognitive functions (12, 54). AD is a devastating cognitive disease accompanied by impairments in perceptive functions including visuospatial abilities and cognitive functions such as memory, executive, and language functions (55). Thus, the altered neocortical hierarchy in patients with AD may be related to the impairment of multiple cognitive functions.

### Gradient dispersion and functional segregation

To be more intuitive, we quantified the functional segregation using gradient dispersion in the gradient space. At both the whole-brain level and module level, AD showed overall decreased dispersion. We applied well-established measures of system segregation and the participation coefficient to further correlate gradient dispersion to the concept of functional segregation (49-51). Consistent with our hypothesis, gradient dispersion was positively correlated with system segregation and negatively correlated with the participation coefficient. These findings indicated decreased segregation in AD. The pathological markers of AD are the presence of amyloid β protein (Aβ) deposition (neuritic plaques, NP) and tau pathology (neurofibrillary tangles, NFT). According to the proposed pathological staging scheme in AD, cortical neurofibrillary pathology first occurs in the transentorhinal and entorhinal regions, then extends to the limbic allocortex and association cortex, and finally influences the secondary and primary regions (56). Patterns of gray matter atrophy in AD fit this pathological staging of AD, first focusing on the medial temporal lobe and then extending to other association regions in the temporal lobe, parietal lobe and frontal lobes (57). As mentioned in the hypothetical model of AD, these association regions overlap with the late-myelinating neocortex regions (including the temporal, parietal, and prefrontal lobes), which are vulnerable to AD pathology (58). Oligodendrocytes produce myelin and cholesterol, and they are especially susceptible in the late-myelinating neocortex regions, in which the myelin of neurons is thinner and easily destroyed (58). With increasing age, oligodendrocyte and myelin breakdown increase iron and cholesterol levels, which can promote the toxicity of the intracortical environment. Increased iron and cholesterol levels may accelerate the oligomerization of Aβ, which inversely impair myelin (58, 59). Myelin breakdown may promote NP- and NFT-related pathological processes(58). These findings indicated that the cortical association regions are vulnerable to pathological impact. In fact, previous studies have shown that in elderly individuals, the association regions overlap with cortical hubs, which are highly connected to many other brain regions (60, 61). Cortical hubs are essential to maintain modular organization and tune brain network organization (62). Higher connectivity between hubs can increase segregation (62). Impairment of hubs leads to global and widespread reorganization of the brain network and decreased segregation (16). Thus, AD-related pathological changes may finally cause decreased segregation. In this study, the overlapping results of changes in LN in the aMCI group may suggest abnormal functional organization in local regions and provide more evidence for the early diagnosis of AD. By correlating gradient dispersion with previously established graph-related measures, our results strengthened the understanding of gradient dispersion. Gradient dispersion, as a more accurate description of the brain regions, showed high sensitivity and demonstrated its advantages as an indicator of functional segregation.

### Cognitive performance and functional segregation

In our study, by combining cognitive performance and gradient dispersion analysis, we demonstrated that decreased functional segregation was related to cognitive decline in aMCI and AD patients. There is some evidence indicating that a higher cognitive performance is related to higher functional segregation (62, 63). Previous studies have demonstrated a trend of decreased segregation with age, which may cause a decline in cognitive performance in healthy elderly individuals (49, 64, 65). In patients with AD, higher cognitive resilience is associated with higher functional segregation (63). Some studies have demonstrated that balanced modular organization in the brain network is crucial for maintaining cognitive function (10, 66-68). In healthy people, hubs have been proven to maintain modular organization (62). In addition, more segregated brains demonstrated higher cognitive functions in multiple cognitive domains (62). By combining the results of previous studies and our results, we inferred that cognitive decline was related to reorganization of the global brain network, which caused abnormal functional segregation. Furthermore, by regarding aMCI and AD as models of cognitive decline, we explored the relationship between cognitive function and functional segregation, which complemented existing studies of the relationship between cognition and large-scale brain networks.

### Methodological considerations

Previous studies on aging have shown that aging is accompanied by changes in the functional modular organization of the brain (43, 44). Functional segregation and specialization are decreased, and the modular structure of the elderly individuals is less distinct, which is different from that of healthy young adults (43, 44). Considering that the participants included in this study were all over 60 years old and that the previous functional atlas based on healthy young adults was not suitable for subjects in this study, we derived a modular structure based on the NC group-level connectome using a modular detection method (45).

Recent studies have demonstrated that the connectome gradient has been used in multiple contexts because of its meaningful representation of brain organization principles. The connectome gradient shows variability among individuals and groups; thus, its reproducibility is important for studies. A previous study suggested that longer time-series data produce more reliable gradients (69). However, it is not practical to conduct long scans in patients with AD. In this study, the length of the time-series data is 8 min, which is a comparatively acceptable length at present (27, 69).

## Limitations

There were some limitations to this study. First, although the NC, aMCI, and AD groups were included in this study, the cross-sectional data may reduce the continuous information in disease progression. At present, we have found that for functional gradient dispersion, patients with aMCI have initial changes in local regions, and patients with AD have macroscale abnormalities in the whole brain. This measure has the potential to be a biomarker for the diagnosis of aMCI and AD. However, the patterns of the continuous changes in disease are still unclear. Longitudinal studies are essential to shed more light on the question of how brain function changes over time in disease populations. In the future, we need to perform dynamic monitoring to investigate the dynamic changes with the progression of the disease in longitudinal studies. Second, the present study only used rs-fMRI to investigate the functional changes in the brain, while combining structure MRI might reveal more features about structural abnormalities and further explain the functional changes. More specifically, this study used functional gradients to capture the functional variance along the cerebral cortex, and we regarded the position of a region in the functional gradient space as its unique functional role (20). Recently, structural gradients based on MRI have revealed gradual transitions of multiple structural features along the cortex, such as microstructural gradients, structural covariance gradients, and morphometric similarity gradients (24, 70, 71). Along the macroscale cortical structural gradient axis, similar positions of the regions occupied represent similar structural features. However, whether changes in macroscopic structural gradient are associated with abnormal structural features in aMCI and AD have not been investigated. In the future, we hope this question could be explored to explain the alteration in functional gradient and provide more evidence to better understand the mechanisms of brain abnormalities in aMCI and AD.

## Conclusions

In conclusion, by applying connectome gradient analysis, we revealed alterations in functional gradients of patients with aMCI and AD compared with NCs. Our findings identified finer topological organization changes and indicated different extents of altered brain network segregation in aMCI and AD, which was related to the level of cognitive performance. These findings provided new evidence for hierarchical alterations in the brain in aMCI and AD patients, which revealed abnormal functional segregation and strengthened our understanding of the functional organization and progressive mechanism of cognitive decline.

## Additional file

Additional file 1: Supplementary Materials. (DOCX)

## Abbreviations

AD: Alzheimer’s disease
aMCI: amnestic mild cognitive impairment
NC: normal control
rs-fMRI: resting-state functional magnetic resonance imaging
MoCA: Montreal Cognitive Assessment
AVLT: Auditory Verbal Learning Test
CDR: Clinical Dementia Rating
HAMD: Hamilton Depression Rating Scale
ADL: Activities of Daily Living
NIA-AA: National Institute of Aging-Alzheimer’s Association
MPRAGE: magnetization-prepared rapid gradient echo
HCP: Human Connectome Project
NMI: normalized mutual information
ANOVA: analysis of variance
FD: frame-wise displacement
FEW: family-wise errors
FDR: false discovery rate
SN: somatomotor network
VN: visual network
VAN: ventral attention network
DAN: dorsal attention network
LN: limbic network
DMN: default mode network
Aβ: amyloid β
protein; NP: neuritic plaques
NFT: neurofibrillary tangles.

## Declarations

### Ethics approval and consent to participate

This study was approved by the Ethics of the Medical Research Ethics Committee in Xuanwu Hospital, and written informed consent was obtained from all participants.

### Consent for publication

Not applicable.

### Availability of data and materials

The data, material, and code that support the findings of this study are available from the authors upon reasonable request and with permission of the corresponding author.

### Authors’ contributions

SL contributed to the study design. YH performed the image data acquisition and behavior data collection. YrH, QL, and ZF performed the data preprocessing, data analysis, and interpretation. YrH, QL, DZ, ZF, and SL wrote the manuscript. All authors reviewed the manuscript.

## Acknowledgements

Funding from Startup Funds for Top-notch Talents at Beijing Normal University and National Natural Science Foundation of China (Grant Nos. 81972160) are gratefully acknowledged.

